# Neural evidence of cognitive conflict during binocular rivalry

**DOI:** 10.1101/2019.12.19.873141

**Authors:** Alice Drew, Mireia Torralba, Manuela Ruzzoli, Luis Morís Fernández, Alba Sabaté, Márta Szabina Pápai, Salvador Soto-Faraco

## Abstract

To make sense of ambiguous and, at times, incomplete sensory input, the brain relies on a process of active interpretation. At any given moment, only one of several possible perceptual outcomes prevails in our conscious experience. Our hypothesis is that the competition between alternative representations induces cognitive conflict, eventually leading to fluctuations between different perceptual interpretations. We used binocular rivalry, a popular protocol to probe changes in perceptual awareness [1–3] and drew on the conflict monitoring theory, which holds that cognitive control is invoked by the detection of conflict during information processing. We looked for an increase in power of fronto-medial theta oscillations (5-7 Hz), an established EEG marker of conflict detection [4–7]. Our results show that fm-theta power increases right before perceptual alternations and decreases thereafter, suggesting that conflict monitoring is related to perceptual competition. Furthermore, to investigate conflict resolution via attentional engagement, as held by the conflict monitoring theory [8], we also looked for changes in parieto-occipital alpha oscillations (8-12 Hz) associated to perceptual switches. These oscillations have been associated to attention allocation via functional inhibition in sensory cortices [9–12]. The power of parieto-occipital alpha was inversely related to that of fm-theta, reflecting periods of high inter-ocular inhibition during stable perception, and low inhibition around moments of perceptual change. Our findings validate a prediction made 20 years ago in the seminal paper formulating the conflict monitoring theory, and establish a previously unknown link between conflict mechanisms and the processes leading to perceptual awareness.

**Highlights:**

- Binocular rivalry induces conflict between competing perceptual representations
- Mid-frontal theta power increases around the resolution of perceptual conflict
- Posterior alpha decreases reflect neural excitability around perceptual switches
- The results link cognitive conflict theory with perceptual inference

## Results and Discussion

Few perceptual phenomena have aroused as much curiosity as binocular rivalry (BR) - and done so consistently over more than 400 years. In the 1970s, it was even employed as an artistic technique when Salvador Dalí used stereoscopic perception in a series of paintings, as part of an ongoing attempt to challenge our perception of reality. BR arises when two disparate images are presented separately to each eye, disrupting binocular fusion and producing stochastic alternations between both. The phenomenon has been of continual interest for the study of visual perception and awareness, as it induces striking fluctuations in perceptual experience despite constant physical stimulation [1–3]. Here, we used BR to test the relationship between visual perception and cognitive conflict, and reasoned that the competition between perceptual interpretations of the same sensory input triggers cognitive control mechanisms. Despite being a predicted outcome of the cognitive control framework [8], the link between perceptual ambiguity and conflict has not yet been established.

Cognitive control refers to a set of functions that allow the configuration of the cognitive system to perform tasks flexibly, according to instructions, internal goals and changes in the environment. Botvinick and colleagues [8,13] proposed that the anterior cingulate cortex (ACC), a fronto-medial (fm) brain area, could be responsible for the detection of conflict during information processing (i.e., when two incompatible representations are simultaneously active) [14–16] as well as for triggering strategic adjustments (i.e., attention orienting) for conflict resolution [8]. Conflict detection is also reflected by increases in oscillatory power within the theta band (5-7 Hz) over frontal electrodes in the EEG [4–7]. Although the conflict monitoring theory [8,13] is firmly grounded in response conflict (i.e., when a prepotent automatic response must be overridden in order to make way for the correct response, as in the Stroop task for example), it acknowledges that conflict may occur at various stages of information processing including perceptual representation and stimulus categorisation (verbatim from Botvinick and colleagues [8] p. 464) *‘If the ACC does respond to stimulus conflicts, one might expect this area to become active under conditions of binocular rivalry or in viewing ambiguous figures’*). However, stimulus conflict, unlike response conflict, remains a little explored phenomenon.

Here, we hypothesised that (1) Stimulus-based conflict would be reflected by an increase in mid-frontal theta power shortly before the moment of conflict resolution occurring during perceptual switches in BR; and that (2) The subsequent involvement of attention is expressed as a decrease in occipital alpha power prior to perceptual switches, reflecting fluctuations in the excitability of parieto-occipital areas. It is important to note that alpha is a physiological marker of attentional allocation via functional inhibition across the sensory cortices. These hypotheses, the analysis pipeline and the experimental procedures described hereafter were pre-registered on the Open Science Framework (https://osf.io/g4hzp/).

In the experiment, participants (n=32) performed a binocular rivalry task while EEG data was recorded. The rival stimuli were Gabor gratings of different colour (red/green) and orientation (±45° tilt), which had been adjusted in luminance for each eye of each individual to achieve approximate equal dominance. Participants experienced rivalry by looking at the stimuli through stereoscope mirror glasses and reported their percepts on a keyboard. Dominance periods had an average duration of 2.34±0.76 s (red percepts), 2.57±0.83 s (green percepts) and 1.88±1.06 s (mixed percepts). The mean number of red percepts was 140.71 ± 43.89 (29.12%) and of green percepts was 159.42 ± 44.09 (35.44%). Importantly, the total percentage of time spent in the ‘Null percept’ was low (9.12 s [0.84%]), which we expected, since the rival stimuli are continuously present.

The main EEG analysis focused on the periods bounded between perceptual switches, termed trials (we selected trials of 1.5 s or longer; average number of trials per participant 142±48). We selected 500 ms time windows right before and right after the reported perceptual switches. Results showed that the power in fm-theta oscillations increased before a perceptual switch and decreased immediately after it (see Figure 1) according to the prediction, namely that theta power would be higher shortly before the reported perceptual switch (the pre-switch window), compared to the post-switch window (t(29)=3.3017, one-tail, α=0.05; p=0.0013; Cohen’s d=0.60). We interpreted fm-theta power changes as a marker of conflict detection resulting from the competition and reciprocal inhibition dynamics between the neural populations governing each percept, in line with current accounts of binocular rivalry [17,18].

**Figure 1:**
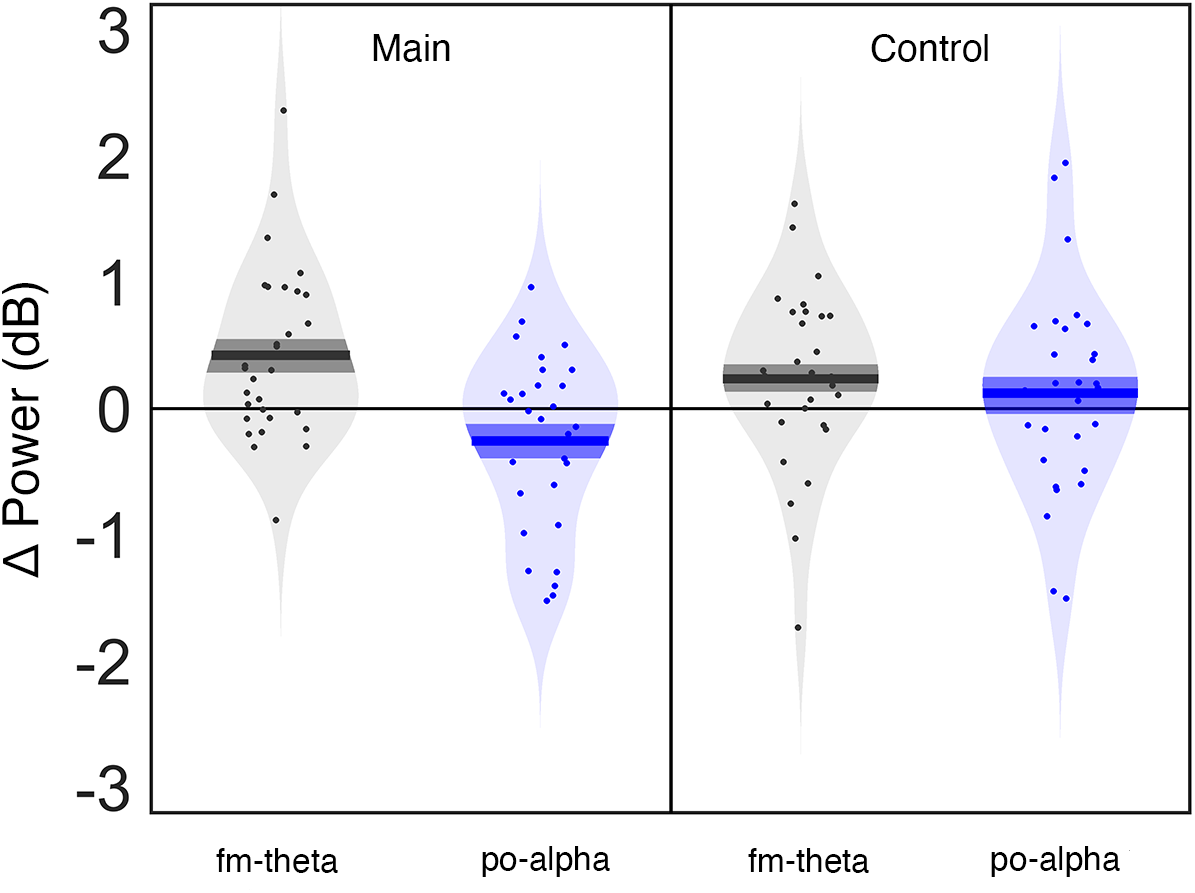
Left: The results for the preregistered analysis presented the expected pattern: Fm-theta power (dB) post-switch was significantly higher than fm-theta power pre-switch (grey), whereas parieto-occipital alpha power (dB) post switch was significantly lower than parieto-occipital alpha power (dB) pre-switch (blue). Right: In the control analysis, after time shifting our windows of interest, fm-theta power (dB) postswitch was still significantly higher than fm-theta power (dB) pre-switch (grey), whereas the effect for parieto-occipital alpha was not observed anymore (blue). Solid lines correspond to the group mean of each of the conditions, dark shaded areas correspond to the mean ± standard error of the mean. Light shaded areas correspond to the distribution of the individual data and dots correspond to individual data.

One point of consensus is that BR is due to one or more mechanisms of reciprocal inhibition between competing neural populations governing each percept at different stages in the visual pathway [18]. The population with the strongest signal determines the dominant percept until a process of neural adaptation diminishes its responsiveness and the alternative competitor population eventually takes over as the dominant percept [17]. We reason that the theta oscillations mirror the dynamics of competition throughout the cycle of conflict monitoring, detection and resolution. We propose that the conflict resulting from competing perceptual representations in BR plays out continuously. When the competition has a strong winner the conflict signal (and therefore theta power) is low; this happens putatively during periods of stable percepts. On the other hand, when the competition is steep there is no strong winner, so conflict is higher (hence, higher theta power); we believe this happens in moments impending a change in percept. Furthermore, the outcome of this competition necessarily fluctuates over time since the conflict in BR can only be temporarily resolved until the inhibitory signal from the neural population governing the competing percept begins to weaken, calling the alternative representation to awareness. Incidentally, the possible role of frontal brain areas seemingly brings new evidence in an age-old debate [19–22] that this competition plays out in the visual pathway according to “top-down” processes [23–25].

The conflict monitoring theory also suggests that one of the functions of the ACC is to trigger strategic adjustments (i.e., attention allocation) for conflict resolution [8]. In the case of BR, attention becomes relevant insofar as there is conflict to be resolved due to sustained visual competition [26]. Furthermore, the dynamics of BR have proved particularly sensitive to attentional modulation, slowing down significantly when attention is directed away from the rivalling stimuli [27–31]. Here, we adopted the ‘sensory gating hypothesis’, which suggests that alpha oscillations reflect attentional selection via functional inhibition of irrelevant information across the sensory cortices [9–12]. For example, alpha decreases have been related to attentional adjustments after errors in a Stroop task [32]. We therefore homed in on occipital alpha-oscillations as a marker of selective inhibition during BR dynamics. Specifically, if BR induces perceptual conflict between stimuli, periods of low inhibition between percepts (low alpha) would co-occur with stronger competition, possibly preceding an impending perceptual switch. Conversely, periods of high inhibition between percepts (high alpha) would co-occur with weak competition, which might immediately follow a perceptual switch. Our analysis revealed that alpha oscillations decreased right before a perceptual switch and increased immediately after it (t(29)=-2.2327, one-tail, α=0.05; p=0.0167; Cohen’s d=-0.40) reflecting evidence in favour of the engagement of attention allocation via inhibition (see Figure 1).

Despite finding statistical evidence of the antagonistic pattern of power modulation in fm-theta and posterior alpha oscillations around moments of perceptual fluctuation, one concern for interpretation is the possibility of motor contamination from the keypress reports in the time windows of interest. Our paradigm entails decision-making, motor planning and motor execution and solving it requires participants to almost constantly be pressing at least one key. Regardless of the use of ROIs (cf. STAR Methods, *Time frequency analysis*), the corresponding scalp topographies (see Figure 2) are not precise as to the origin of the spectral patterns of interest, given the low spatial resolution of EEG. Furthermore, although frontal areas were once thought a prime candidate for spontaneous perceptual changes in multistable perception [33–35], a possible confound between report and introspection-related activity has recently called this into question [22,36,37], with competing interpretations of what frontal activity may represent [38].

**Figure 2:**
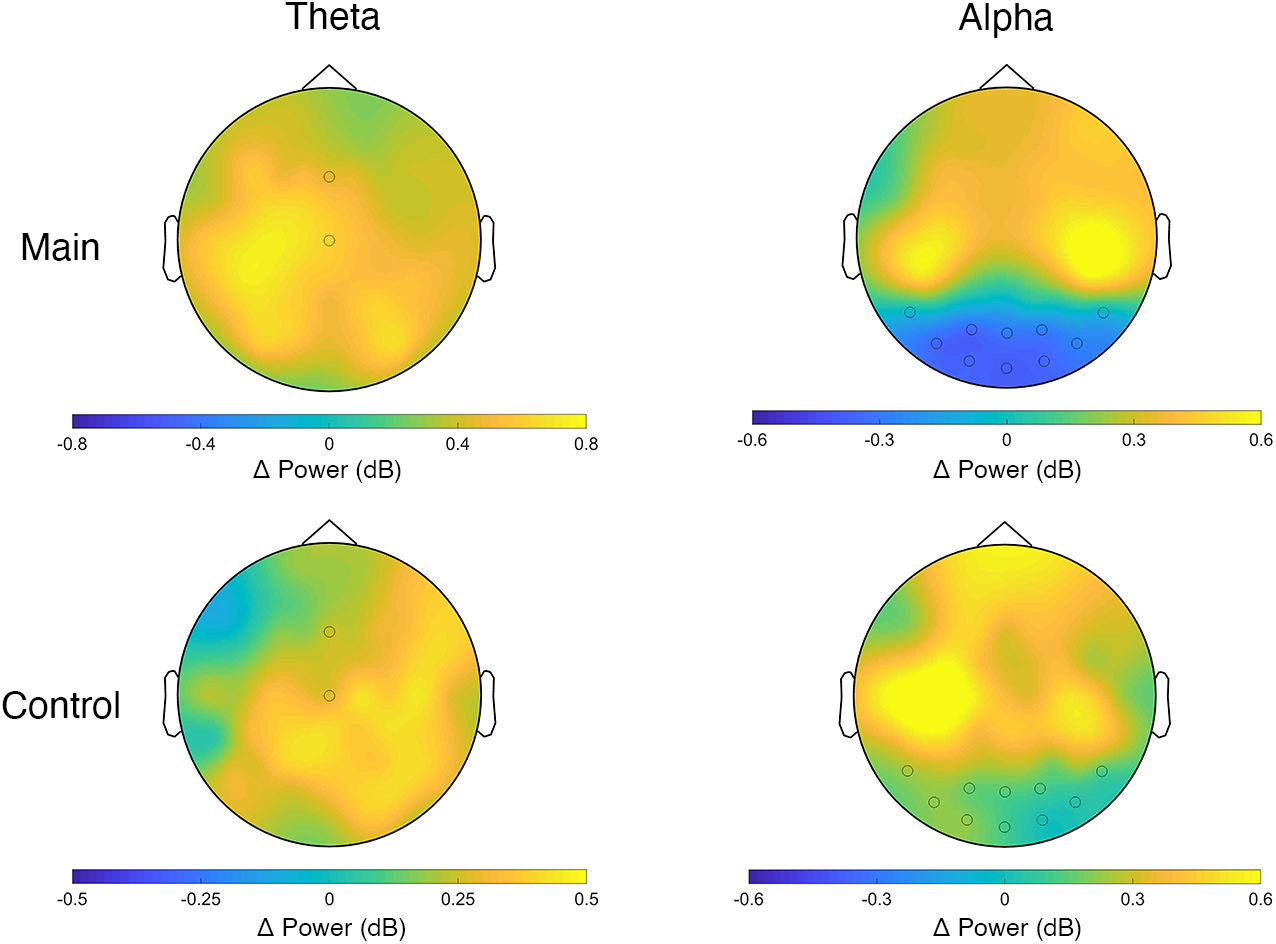
Topographical representation of power contrast (pre switch versus post switch) for the preregistered analysis (top) and control analysis (bottom). Left column corresponds to theta band topographies and right column to alpha band topographies.

To corroborate the robustness of our findings therefore, and to make sure that the changes in theta power were not due to report-related activity, we ran further analyses. We performed a control analysis designed to bypass the period of potential motor contamination by shifting the windows of interest in time, away from the keypresses. To do so we used trials of 2,246 ms or longer (92±40 trials per participant). The pre-switch window was shifted by 446 ms before the switch (the average response time in a replay condition using physical alternation instead of binocular rivalry, cf: STAR Methods). The post-switch window was shifted by 300 ms after the response, which was the estimated duration of the motor evoked potential [39]. As in the main analysis, the fm-theta power modulation was significant (t(28)=1.9809; p=0.029; Cohen’s d=0.37) with shifted time windows, however no reliable effect was seen in alpha oscillations: (t(28) =0.79; p=0.8; Cohen’s d=0.16) (see Figure 1).

**Figure 3:**
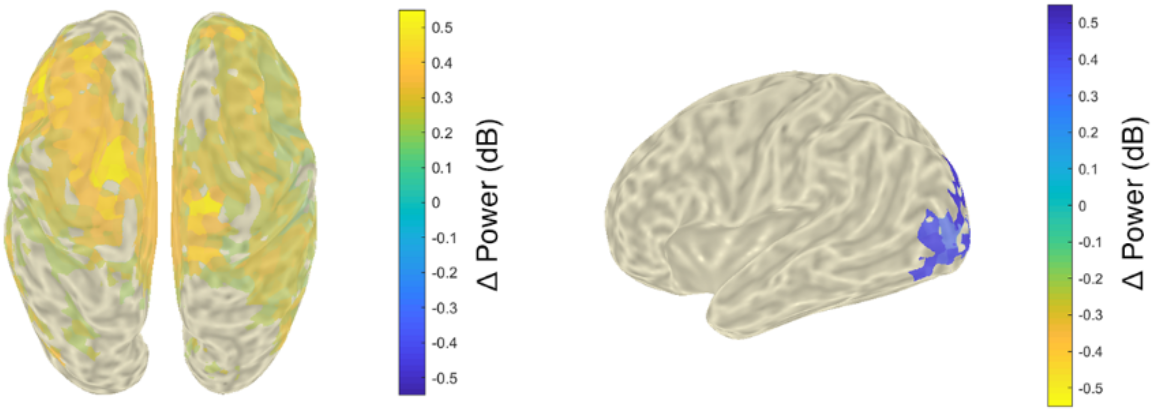
Brain topographies show source localised activity for theta (left) and alpha (right) pre-post switch contrasts.

The results of this control analysis help separate fm-theta changes from responses, but leave open the possibility that alpha modulations are conflated with response-related factors. This is because, in the main analysis performed in near vicinity of perceptual switches, both the response-conflict and the perceptual-conflict interpretations make indistinguishable predictions regarding alpha activity at the sensor-level. That is, strategic adjustments through attentional modulation via inhibitory processes may only be present in the immediate temporal vicinity of the perceptual switch, thus undetectable when shifting the windows of interest in time. Indeed, it has been suggested that the mechanisms for cognitive control operate according to a temporal hierarchy [40]. In order to rule out contamination by motor report in the vicinity of perceptual switches, we performed a source localisation of the theta and alpha modulations found in the main analysis (see Figure 3). We used dynamic imaging of coherent sources (DICS), a spatial filtering technique operating in the frequency domain [41] (see supplementary materials and methods for details). As expected, theta power modulation had a clear source consistent with fronto-medial areas, hence the predicted ACC generator. Therefore, the increase in theta oscillatory power did not arise from motor areas due to response, but can rather be attributed to perceptual conflict. Importantly, the source of alpha activity was localised in posterior occipital regions consistent with visual areas, clearly away from motor-related areas.

According to our hypothesis, the time course of theta and alpha oscillatory power should present opposite patterns during the duration of a percept. In order to appreciate this, we explored the temporal evolution of theta and alpha activity. We selected trials with long percepts (the same ones used for the control analysis) and calculated oscillatory power using 500 ms sliding windows (steps of 2 ms) in the corresponding ROIs for the theta and alpha frequency bands. For each percept, the power was normalised with respect to the first 500 ms right after the key press (in dB). Then, for each trial, we selected the windows centred at −20% to 120% (in steps of 10%) of the total duration of the percept. Finally, we calculated the theta and alpha time courses for each percept and participant. The resulting averaged time courses (represented in Figure 4. A.) illustrate the complementary patterns of frontal theta and posterior alpha. This pattern can be observed on a trial-by-trial basis, in Figure 4. B. (trials ordered by duration). Altogether, the complementary time courses of frontal theta and posterior alpha give credibility to the interplay between conflict mechanisms and perceptual processes.

**Figure 4:**
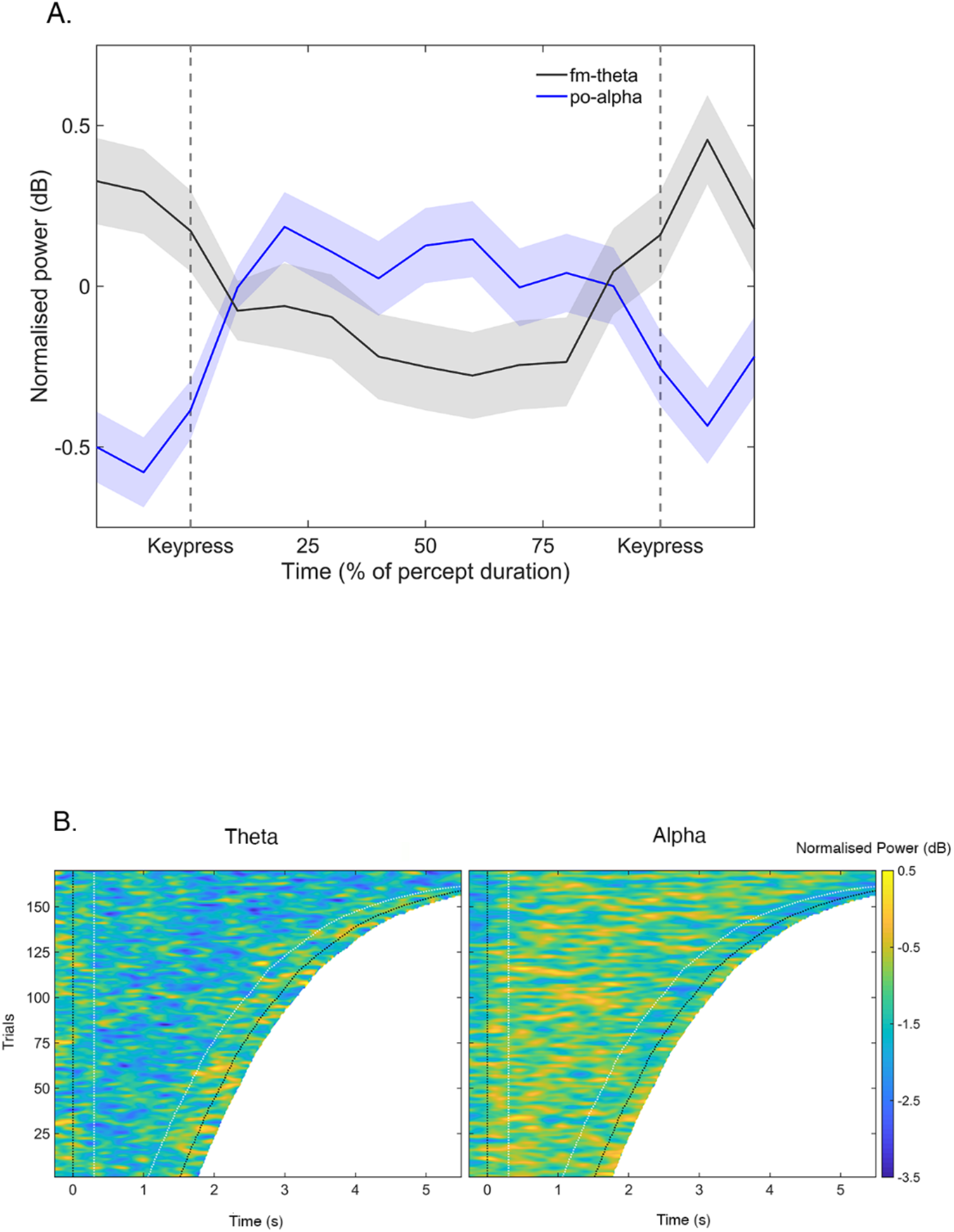
**A.** Timecourse of fm-theta (grey) and posterior alpha (blue) oscillatory power in the period between keypresses denoting perceptual switches, averaged across all trials of all participants. The activity has been measured in steps of 10% of total percept duration. **B.** Left: Temporal evolution of theta power (normalised with respect to mean power during percept) for trials of all participants sorted from shortest to longest. Right: Temporal evolution of alpha power (normalised with respect to mean power during percept) for trials of all participants sorted from shortest to longest. Black lines correspond to key presses and white lines to estimated ERP and RT.

Based on the findings reported here, we believe that the theory around conflict may fit well with predominant frameworks of perception, such as models based on predictive coding [42]. According to these models of perception, the brain relies on active interpretation (inference) in order to make sense of the vast amount of information it has to deal with. Ongoing inferences are constantly adjusted as perceptual predictions are compared against incoming sensory input. We propose that when perceptual predictions mismatch incoming input, conflict is detected and the necessary adjustments are invoked in order to stifle it. That is, changes in the oscillatory marker for conflict (theta) seem to be entangled with complementary changes reflecting the attentional allocation (alpha), during perception. Furthermore, theta and alpha-band oscillations have the intriguing characteristic of presenting inverse patterns in different brain areas despite fluctuating at neighbouring frequencies (respectively 5-7 and 8-12 Hz). Although we were unable to preclude competing interpretations for our findings in the alpha band based on the sensor-level results alone, a posteriori source localisation pointed to the expected involvement of posterior parieto-occipital alpha oscillations, inversely related to fm-theta oscillations. Future research will help relate these complementary oscillatory patterns to the unifying framework of conflict detection and resolution in perception.

In sum, we provide supporting evidence of cognitive conflict occurring between rival stimuli during BR. These findings provide a step towards a broader understanding of the role of cognitive conflict in perception. Should the role of conflict become generalised beyond these initial results, it will provide a promising way of understanding perception within and across sensory modalities, as well as perceptual decision-making.

## STAR Materials and Methods

### Participants

Thirty-two healthy participants (15 female, aged between 18-34 years, mean age of 23) participated in the experiment. Although our pre-registered idea was to record 25 participants, we recorded 32 as this was the required sample size for a parallel study, using the same experimental design and participant data. Inclusion criteria comprised normal or corrected to normal vision and no medication. Participants received 10€/ hour in return for their participation. They all provided written informed consent prior to the study and were naive to the purpose of the experiment. One participant was discarded before starting the experiment due to line noise in the EEG recording. The study was run in accordance with the Declaration of Helsinki and the experimental protocol approved by the local ethics committee in Parc de Salut Mar (Universitat Pompeu Fabra, Barcelona, Spain).

### Apparatus and Stimuli

Participants were presented with two static circular rival Gabor patches (11.5° diameter; 0.1 contrast and 0.73 cycles/cm of spatial frequency and gratings oriented at ±45°) and different colours: red and green. Images were presented on a 19.8-inch CRT monitor (1024×768; 120 Hz refresh rate) with a grey background (10.7 cd/m^2^) displayed at 80cm from the participants’ eyes. Each image was presented to a different eye by means of stereoscope mirror glasses. Visual stimuli were created using the MATLAB Psychtoolbox toolbox (version 3.0.12 (adaptive procedure from Pelli [43]). and Matlab version R2017b [9.3.0.713579]). These parameters gave us a binocular rivalry paradigm in which percepts can last longer than 1 second when active behavioural report is requested from the participants.

The green RGB value was matched to the subjective luminance of the red Gabor using an up/down procedure adapted from Cavanagh and colleagues [44], in order to minimise the flickering between red and green RGB Gabors at a rate of 60 Hz. To achieve this, participants were instructed to regulate the flickering until it had stopped completely, indicating that the colours had been matched. This served to prevent potential dominance of one percept based on stimulus features [18] (cf. *Results* section: *Behavioural*).

### Procedure

Prior to the experiment, participants were dark-adapted for 5 minutes. The stereoscope mirrors were calibrated for stimuli to appear at the same retinal location of each eye, by ensuring monocular vision. Subjective luminance of stimuli was matched.

During the BR blocks, stimuli were presented continuously throughout a block. Since the rival Gabors had dissimilar colours (i.e., red and green) and grating orientation (±45° tilt), observers experienced rivalry as perception alternated between the two gratings.

Participants were seated in a dimly lit and sound-attenuated room and asked to keep movements (including eye movements and blinks) minimal during each block. The experiment consisted of 13 blocks lasting 120 s each. Before the experiment, participants performed two training blocks in order to familiarise with the task (training data were not analysed). Experimental blocks were interleaved with breaks (the duration of which was self-paced by participants) to limit fatigue. Unbeknownst to them, the task was divided into two conditions: 9 blocks of a binocular rivalry and 4 blocks of a replay condition (see Results and Discussion), with one replay block following two consecutive rivalry blocks (in an AAB alternation pattern, plus one additional rivalry block at the end). As our study mainly targeted rivalry blocks, we included the replay condition only in order to have an estimate of the ability to report alternations in the BR blocks and the latency of this report. In the replay condition, the same image was presented to both eyes, and we recreated the perceptual fluctuations, akin to those in the BR condition, by physically alternating between the images. The temporal dynamics of the fluctuations were determined individually by the participants’ cumulative distribution of percept-dominance from all previous rivalry blocks. The mean and standard deviation of percepts (mixed, green and red) were used to generate three gamma distributions of percept durations from which the simulated percepts durations were drawn. Importantly, the total percentage of time spent in the ‘Null percept’ during replay was low, (1.89 s [0.39%]) indicating a behavioural performance in line with the expectation.

Participants were instructed to fixate their gaze on the central fixation cross. The keys used for the perceptual report were X and D on a QWERTY keyboard which participants pressed with the left and right index fingers (i.e. continuously pressing X, D or both keys to report one colour, the other or a mixture, respectively). The assigned key for the green and red stimulus percepts was counter-balanced across participants. To report a mixed (piecemeal) percept, participants were instructed to press both keys simultaneously. In order to report none of the three aforementioned percepts (null percept), they were instructed to depress both keys. The eye (left/right) in which each stimulus was physically presented was also randomised across participants in order to avoid biases due to eye dominance (red left/green right; green left/ red right).

### EEG recording

During the experiment, electroencephalographic (EEG) data were acquired using a 60 electrode system (actiCAP, Brain Products GmbH, Munich, Germany) placed in accordance with the 10-10 international system. The ground electrode was placed on AFz and the online referenced on the tip of the nose. Electrodes for offline re-reference were placed on right and left mastoids. The vertical electrooculogram (Veog) was recorded by an electrode underneath the right eye, and the horizontal electrooculogram (Heog) at the outer canthus of the right eye. Impedance was kept below 10kΩ for all electrodes. The signal was recorded via BrainVision Recorder (Brain Products GmbH, Munich, Germany) at a sampling rate of 500 Hz.

### EEG Pre-processing

Pre-processing and analyses of EEG data were done using Fieldtrip [45] and custom-made code in Matlab. Data were separated into segments corresponding to perceptual switches, i.e. occurring between keypresses. Switches from pure percept (red or green) to another pure or mixed percept were included. Data were inspected for artefacts in order to manually reject segments contaminated by blinks, movements and noise upon visual inspection. Participants with fewer than 30 trials in at least one of the conditions of interest were discarded from analyses. One participant was discarded based on the criterion for minimum number of trials in the main analysis, whereas two participants were discarded in the control analysis, resulting in 30 subjects for the main analysis and 29 in the control analysis (see below).

### Behavioural analysis

Percepts shorter than 300 ms (i.e. a rough estimation of the mean latency of motor evoked potential from human movement such as keypresses [39]) were discarded. In order to check for dominance-bias and reliability of participants’ report, we extracted for each participant the mean lengths of percepts, the total and percentage of time spent in the null percept condition during rivalry and replay conditions, and the mean percentage of time spent in each percept. Evidence in favour of reliable report and against a bias towards one of the stimuli should be reflected by a similar duration of dominance for red and green percepts, due to the subjective isoluminance adjustment. Evidence in favour of the reliability of participants’ report should be reflected by a low percentage of time spent in the null percept in both conditions (cf. Results and Discussion). This is because in the rivalry condition stimuli were presented continuously to the eyes, whereas in the replay condition we did not include null percepts, therefore correct reports should match the physical alternations presented. Given our efforts at tuning stimuli parameters in order to match the strength of the stimuli, we did not expect the duration of mixed percept to exceed that of pure percepts. Based on our results (cf. Results and Discussion), no participant was discarded based on behavioural performance.

### Time-frequency analysis

According to the pre-registered hypotheses (https://osf.io/g4hzp/), the analysis was centred at two frequency bands: theta (5-7 Hz) and alpha (8-12 Hz) in two 500 ms time windows. Only segments that were free from artefacts and had a minimum duration of 1.5 s were included in the registered analysis. This segment length ensured the time necessary for analyses of the intended time windows, at the intended frequency bands. Window length was selected in order to include at least 3 cycles of the central frequency (6 Hz) in the slowest (theta) frequency band of interest. One time window was located at the beginning of the segment, right after a keypress (henceforth referred to as post-switch), and the other was located at the end of the segment, just before a keypress (henceforth referred to as pre-switch). We used a Fast Fourier transform with a Hanning taper to extract the power in the frequency bands of interest in the pre- and post-switch windows, zero padded up to a length of 1 second.

The measure of interest was the power in the pre-switch versus post-switch, measured in decibels (dB). Activity was measured over the whole scalp, but regions of interest were pre-defined for each frequency band: a fronto-medial region (Fz, Cz) for the theta band and a parieto-occipital region (P7, P8, PO7, PO3, POz, PO4, PO8, O1, Oz, O2) for the alpha band. The power contrast was calculated for each frequency and electrode of interest and, subsequently, averaged across frequencies and ROIs. Mistakenly (and noted a posteriori) data from FCz was not recorded during the experiment, therefore data was acquired by pooling from Fz and Cz given the spatial resolution.

A control analysis was performed trying to estimate the contamination from motor evoked potentials in the post-switch window and from activity due to active report (decision-making, motor preparation and motor execution of the keypress) in the pre-switch window. The response time of participants when reporting perceptual transitions (excluding mixed and null percepts) was estimated from the response latency in the replay condition (which resembled a visual detection task and it was included specifically for this purpose). The median response time across participants in the replay condition was 446 ms (SD=103 ms). In the control analysis, then, the pre-switch window was shifted by 446 ms, while the post-switch window was shifted by 300 ms, corresponding to the duration of a motor-evoked potential from human movement such as a keypress [39].

### Source localisation

The leadfield was calculated using a standard boundary element method available from Fieldtrip, version 20190203, the grid spacing was set to 10 mm [45]. For each participant, frequency band (theta and alpha) and condition (Registered and Control), power and cross-spectral density were calculated at the pre-switch and post-switch time windows with the same parameters as in the electrode level analysis. For each participant, frequency band and condition, we calculated the common filter for pre-switch and post-switch time windows using DICS, with 1% regularisation. Then, we used the common filter to obtain the power at the pre-switch and post-switch time windows for each frequency band and condition and we computed the power differences (Pre-switch vs Post switch) in dB for each participant, frequency band and condition. Significant pre-switch vs. post switch window power differences at source level were assessed by means of one tailed paired t-tests for each frequency band and condition separately. Multiple comparison correction was performed by means of false detection rate (fdr) [46] procedure at an alpha level of 0.05.

## Acknowledgments

This research was supported by the *Ministerio de Economia y Competitividad* (PSI2016-75558-P AEI/FEDER) and AGAUR *Generalitat de Catalunya* (2017 SGR 1545) grants to SSF. MR is supported by the European Commission Individual Fellowship (Ctrl Code – 794649, H2020-MSCA-IF-2017). AD is supported by AGAUR *Generalitat de Catalunya* FI-DGR 2019.

## Author Contributions

A.D. contributed to experimental design, collected data, contributed to data analysis and interpretation and wrote the paper. M.T. designed the experiment, programmed experimental protocol, analysed and interpreted the data, and wrote the paper. M.R. designed the experiment, interpreted the data, and wrote the paper. L.M.F. designed the experiment, programmed experimental protocol, analysed and interpreted the data and wrote the paper. A.S. contributed to experimental design, collected data, collaborated in data analysis and interpretation. M.S.P. designed the experiment, collected the data, interpreted the data. S.S.F. designed the experiment, analysed and interpreted data and wrote the paper.

